# Whole-body central processing of lateral line inputs encodes flow direction relative to the center-of-mass

**DOI:** 10.1101/2025.06.16.659910

**Authors:** Elias T Lunsford, Martin Carbo-Tano, Claire Wyart

## Abstract

From shifting visual scenes to tactile deformations and fluid motion, animals must interpret patterns of sensory flow around their body to construct stable internal models and produce adaptive behavior. Understanding of how such transformations are encoded within the brain remains incomplete. To tackle this question, we leverage the lateral line of larval zebrafish as a tractable sensory system sensitive to fluid motion that is used to steer navigation, feed, and avoid predators. By presenting stimuli of either direction to neuromasts along the body, we used high-resolution calcium imaging to map hindbrain responses. Unexpectedly, our findings challenge the notion that central lateral line processing lacks topographic structure by revealing a simple yet powerful principle centered on a egocentric spatial framework: the direction and location of local flow motions are encoded in reference to the animal’s center-of-mass. This simple representation enables the brain to register complex flow patterns and provides a robust basis for subsequent behavioral action selection. MON neurons that encode flow toward the center-of-mass broadly project to form bilateral connections onto reticulospinal neurons that coordinate forward locomotion while MON neurons that encode flow away from the center-of-mass displayed a more selective and unilateral projection profile to command neurons for turns. Our discovery represents a shift from purely somatotopic encoding toward an integrative representation of axial position and directionality combined, revealing a novel principle of encoding spatio-directional cues in the hindbrain. This study advances our understanding of how complex mechanosensory inputs select appropriate motor outputs via simple egocentric neural maps in the hindbrain.

**Significance Statement:** Spatial and directional cues are essential to select appropriate actions in response to changes in the environment. How information from broadly distributed mechanosensors across the entire body occurs in the brain enables motor selection remains elusive. By directionally stimulating each and virtually all neuromasts distributed along the lateral line of larval zebrafish, our study uncovers that spatial and directional inputs from each flow sensors is encoded in the medial octavolateralis nuclei relative to the animal’s center-of-mass in order to subsequently recruit reticulospinal neurons driving forward and turn bouts. These results establish a new framework for understanding how broadly distributed inputs get integrated to recruit motor command neurons responsible for producing diverse behaviors.

## Introduction

Animals must continuously transform diverse sensory inputs into coherent internal representations of the environment to generate appropriate behavioral responses. To guide survival behaviors such as navigation, feeding, and escape, sensory systems must accurately encode both the location and direction of external stimuli. At the population level, spatially distributed sensors have the ability to disambiguate the location and direction of a stimulus but individual sensory afferent fibers can be easily confounded or remain agnostic to the location and direction of the sensory cue. Integration centers must be efficiently organized within the brain to ease sensory processing and subsequent motor decisions. To integrate the key features relevant for guiding behavior, we must resolve how sensory systems encoding dynamic inputs from sensors distributed across the body are compactly represented within the brain.

One common strategy employed by a diversity of sensory systems is the formation of topographic maps^1^. Such maps are well-characterized in systems with compact sensory epithelia. In the visual system, retinotopic maps preserve the continuous spatial relationships of photoreceptors in the retina^2^. Similarly, in the auditory system, tonotopic maps organize cochlear hair cell positions along the continuum of sound frequencies^3^. Other sensory systems integrate discrete stimulus features, such as interaural time and intensity differences, into spatially distinct maps^4,5^. The rodent somatosensory cortex incorporates both continuous and discrete mapping in the barrel cortex providing precise spatial encoding of whisker stimulation^6,7^. Despite these insights for organization in compact sensory organs, how spatial and directional cues from body-wide sensor arrays are processed and represented in the brain remains incompletely understood. One technical challenge lies in selectively stimulating individual sensors, often overlapping in space and receptive field, at the scale of the entire body plan while comprehensively recording the sensory inputs and interneuron responses in the brain.

To tackle this question, we leverage here the lateral line system (LL) of larval zebrafish, which offers a tractable sensory organ distributed along the entire body platform sensitive to fluid motion that mediates essential behaviors such as navigation^8–10^, prey detection^11–13^, and predator avoidance^14,15^. Spatial and directional cues from the fluid environment are essential to accurate action selection of an appropriate motor response^10,15,16^. To effectively sample the location and orientation of the flow, the LL is composed of broadly distributed neuromasts (NMs), comprised of polarized mechanoreceptive hair cells, which are embedded on the head (anterior lateral line; aLL) and body (posterior lateral line; pLL) of the animal to detect and transmit information about changes in flow direction to the hindbrain^17–21^. Individual sensory afferents preserve directional cues at the level of the periphery by only innervating hair cells sensitive to the same flow direction ostensibly leading to preservation of directional cues at the primary projection locus^22–24^. By revealing how spatial and directional information across the entire body is processed in the brain, we aim here to gain fundamental insights into how multidimensional inputs are integrated to subsequently modulate the activity of motor commands driving locomotion.

The primary projection site of the LL is the medial octavolateralis nuclei (MON) located in the rostral-dorsal hindbrain. The MON offers a unique opportunity to dissect causal mechanisms that inform central processing in the vertebrate brain during sensory perception. Across fish phylogenies, anatomical studies have shown that the aLL and pLL nerves terminate onto distinct MON regions, preserving an incipient somatotopic organization^25,26^, but at the physiological level we have yet to uncover any topographical representation of sensor position or directional cues in zebrafish^18,27,28^. To address this gap, we aim here to present stimuli of opposing polarities onto nearly all and each NM innervating the head, body, and tail. This approach provides the most comprehensive map to date of MON responses to spatially distributed stimuli across body axes and directions. We combined precise bi-directional stimulation with high-resolution calcium imaging in the MON to reveal a simple yet powerful organizing principle: directional inputs are encoded in reference to the animal’s center-of-mass, establishing an egocentric spatial framework and challenging the notion that central LL processing lacks topographic structure. This representation enables the brain to register complex flow patterns as motion toward or away from the center-of-mass, providing a robust basis for subsequent behavioral action selection. By elucidating how spatial and directional cues from a distributed sensory array are mapped in the brain, our work advances understanding of how vertebrate systems integrate multi-dimensional inputs to drive context-appropriate motor outputs and may offer a generalizable model for how sensory integration and motor control are coordinated through structured neural maps.

## Results

### Functional identification of MON in larval zebrafish

To functionally define the MON loci in 4-6 days post fertilization (dpf) larval zebrafish, we deployed controlled flow along the anterior-posterior axis to stimulate individual NMs while simultaneously recording neuronal responses in the hindbrain (**Figure 1A**). The anatomical perimeter of the larval zebrafish MON has been originally defined by the primary projection patterns of aLL and pLL afferents^17,29^ whereas the functional space has been confined to stimulation of tail NMs^18,27,28^. To maintain access to NMs from head to tail, we secured paralyzed larvae to the base of a silicone semi-circle using tungsten pins, avoiding the encapsulation of head neuromasts associated with traditional agarose head-restraint (**Figure 1A**). We stimulated individual NMs three times for five seconds^18,30^ with 40 seconds of inter-trial intervals using a gentle unilateral single-pulse flow ejected from a large-tip (∼10 µm) micropipette. In this protocol, we monitored for cupulae and kinocilia deflection along the intended axis (**Figure 1B**) and measured kinocilia height (18.8 ± 3.6 µm), tip-displacement (13.1 ± 4.7 µm), and deflection angle (47.8 ± 9.3°; **Figure 1C**), which overlapped with displacement distances (∼8 - 14 µm) sufficient to elicit the entire range of evoked afferent firing rates^30–32^. Responses in the hindbrain were simultaneously recorded by measuring, typically in 10 planes (z = 100 µm) to capture the entire MON, calcium signals using the pan-neuronal, nuclear-tagged calcium indicator *Tg(elavl3:H2B-GCaMP6f)^jf7Tg^* transgenic line^33^; **Figure 1A**).

**Figure 1.**
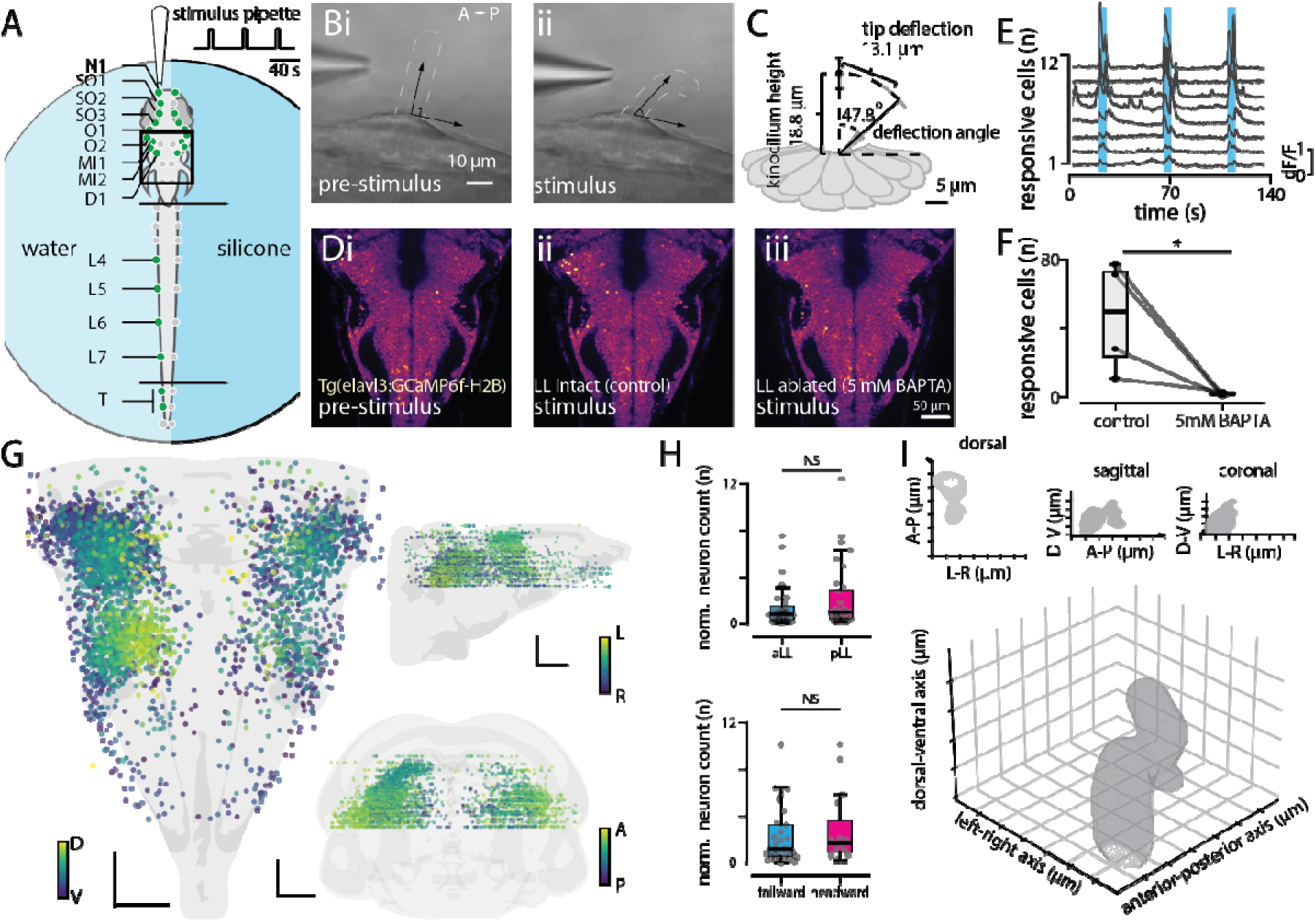
Hindbrain response properties to single neuromast stimulation with unidirectional flow. **A.** Schematic of experimental preparation of a paralyzed larval zebrafish (4-6 dpf) to enable stimulation of individual neuromasts (NM, green) along the head and tail. Black box outlines the field of view of hindbrain imaging (see D). **B.** Brightfield imaging of a single NM before (**i**) and during (**ii**) flow stimulus. Dashed line outlines the perimeter of the cupula and black arrows highlight the angle of the kinocilia deflection. **C.** Average kinocilia height (n = 53, 18.8 ± 3.6 µm), tip displacement (13.1 ± 4.7 µm) and deflection angle (47.8 ± 9.3°) across NMs. **D.** Hindbrain imaging of pan-neuronal, nuclear-targeted genetically encoded calcium indicator in *Tg(elavl3:H2B-GCaMP6f)* transgenic larvae before stimulation (**i**), during stimulation with an intact LL (**ii**) and after chemical ablation of the LL (5 mM BAPTA, **iii**). **E.** Calcium transients from **Dii** of hindbrain neurons that reliably respond during all three stimulus periods (blue bars represent stimulus period). **F.** Box-and-whisker plot representing flow responsive neuron counts before and after BAPTA exposure demonstrates a significant reduction in flow responsive neurons after LL ablation (n = 4 fish / 4 NMs, p = 0.03). **G.** Reference atlas of all flow responsive hindbrain neurons (n = 3,918) across fish (n = 36) and NMs (n = 100) depicted in dorsal (left), sagittal (top-right), and coronal views (bottom-right) color-coded according to depth-of-view. Scale bars: 50 µm. **H.** Box-and-whisker plots of responsive neuron counts normalized by number of fish does not significantly differ when stimulating aLL (39.97 ± 48.89 neurons) versus pLL NMs (51.39 ± 67.43 neurons; Mann-Whitney U stat = 649.5, p = 0.72; top) nor when stimulating with tailward (30.29 ± 34.62 neurons) versus headward flow (28.30 ± 32.33 neurons; Mann-Whitney U stat = 977.0, p = 0.80; bottom). **I.** Three-dimensional mask of the left medial octavolateralis nucleus (MON) generated from contour maps of the 95th percentile from kernel density estimates (KDE) of coordinates from ipsilateral NM connected neurons in the atlas.

Although the MON primarily integrates LL inputs, auditory and vestibular fibers may terminate within the nuclei^34^. To verify that our stimuli selectively targeted the LL, we recorded hindbrain responses to NM stimulation before and after bath application of the calcium chelator BAPTA at 5 mM that disrupts tip-links between stereocilia necessary for mechanotransduction in superficial hair cells^35^ (**Figure 1Di-iii**). Flow-evoked responses (**Figure 1E**) were significantly reduced following this chemical ablation, confirming that the observed hindbrain activity originated from LL input (Shapiro-Wilk test: *p* = 0.372; paired t-test: *t*_(3)_ = 2.75, *p* = 0.035; **Figure 1F**).

Across all 572 trials with consistent recruitment, we identified over 3,900 flow responsive neurons across unique NMs (n = 100) and individuals (n = 36). We registered these flow responsive neurons to a reference atlas (**Figure 1G**) to enable comparisons of response profiles across different stimulus parameters and individuals. On average, stimulation of a single NM led to consistent activation of ∼32 neurons across the entire imaging volume. When comparing the amount of flow responsive neurons when stimulating aLL and pLL or using anterior-to-posterior (tailward) flow or posterior-to-anterior (headward) flow we found no significant difference (p = 0.79, p = 0.80, respectively; **Figure 1H**) indicating no bias in sensitivity to either LL subtype or flow direction when presented with a controlled stimulus. To define the spatial structure of the left MON, we generated a three-dimensional mask by applying kernel density estimation (KDE) at the 95th percentile threshold to the spatial distribution of neurons responsive to ipsilateral NM stimulation (**Figure 1I**). This mask reveals distinct structural patterns, highlighting regions of concentrated flow responsiveness neurons that are not randomly distributed but instead form localized clusters, which may correspond to functional subdivisions within the MON. Altogether, our map provides the most comprehensive representation of the MON in larval zebrafish to date, integrating both aLL and pLL inputs while selectively stimulating individual NMs with directional specificity. This dataset enables a detailed exploration of the somatotopic organization and central processing mechanisms underlying whole-body sensory integration in a vertebrate species.

### Sensory somatotopy within MON represents sensor position across body axes

Lateral line afferents lack contralateral projections and maintain lateralized separation of left and right prior to interfacing with the hindbrain^17^, which may influence how the MON processes left-right asymmetries in flow-evoked signals. To verify our method and investigate the recruitment of MON neurons during single-NM stimulation across the left-right axis, we bootstrapped random samples of NMs located on either the left or right side of the head and calculated the relative density of tailward flow-evoked responses in the hindbrain (**Figure 2**). We found significantly more flow-evoked responses ipsilateral to the stimulated NMs in the prepontine, pontine, and retropontine (r1-6) of the hindbrain for both left and right NMs (p < 0.001; **Figure 2A-C**). While most MON responses were ipsilateral, a subset (∼18%) exhibited contralateral activation (**Figure 2Biii, Ciii**). These findings verify that flow-responsive neurons in the MON predominantly encode lateralized inputs, preserving spatial segregation of sensory signals necessary for behavioral algorithms that facilitate rheotaxis^10^. The LL system extends well beyond the head, raising the question of whether a similar somatotopic organization exists along the anterior-posterior axis, where spatial encoding could support directional flow processing and sensorimotor integration.

**Figure 2.**
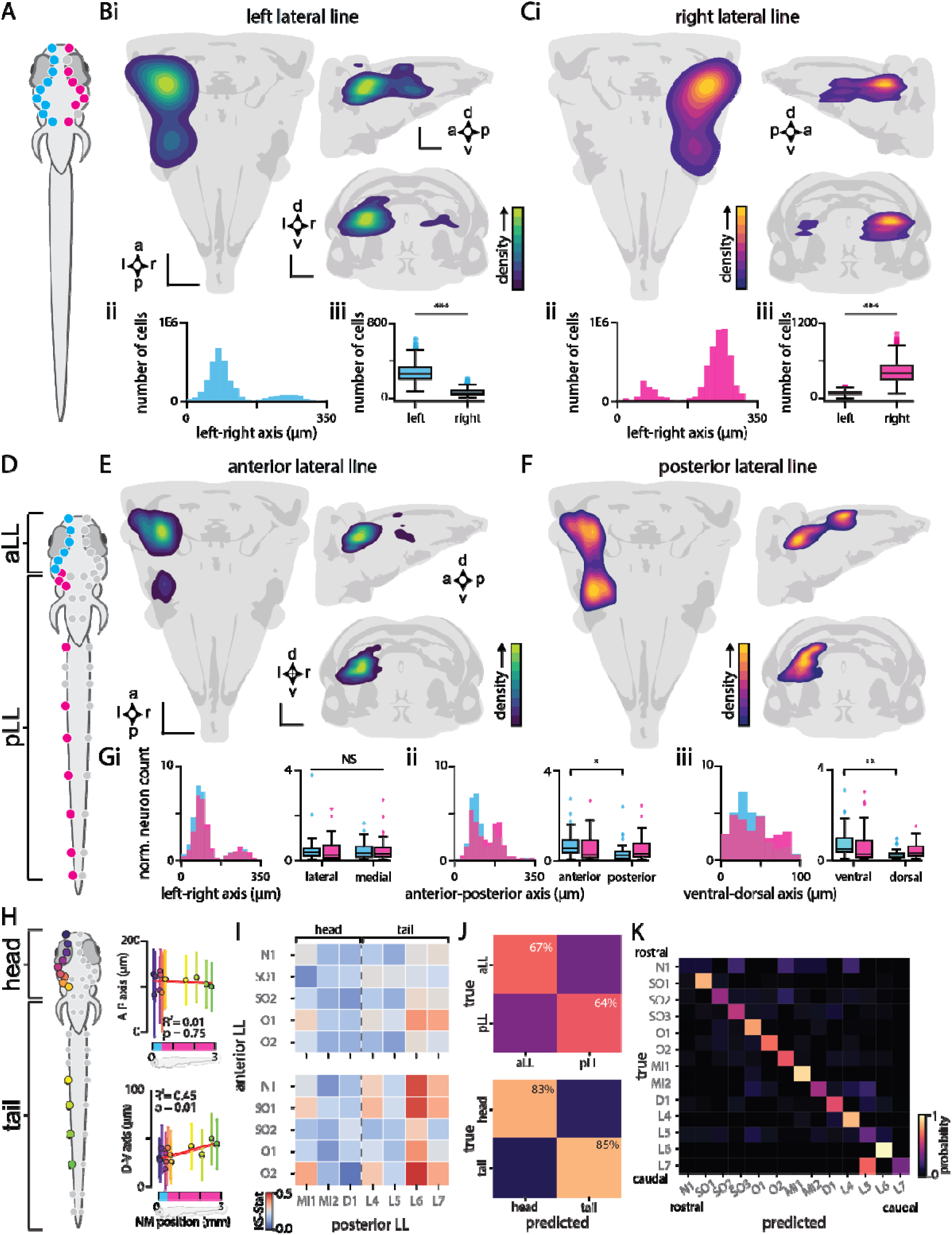
Flow responsive MON neurons demonstrate ipsilateral somatotopy and reliably predict neuromast position along the rostral-caudal axis. **A**. An illustration of a larval zebrafish indicating the position of individually stimulated neuromasts (NMs) located on the left (cyan) or right (magenta) side of the head. **Bi**. Contour heat maps in dorsal, sagittal, and coronal views represent the relative kernel density estimates (KDE) of flow responsive hindbrain neurons that responded when stimulating individual NMs (randomly sampled n = 10, bootstrapped n = 1k) on the left-side of the head. **ii** Histogram visualizes distribution of responsive neurons along the left-right axis. iii. Quantification of significantly more responsive neurons ipsilateral to NMs stimulated on the left side of the body represented in a box-and-whisker plot (N = 29 fish, n = 54 NMs, n = 1,799 neurons; Mann-Whitney U-stat = 99x106, p < 0.001; **iii**). **Ci-iii** A mirrored pattern of ipsilateral activation along the left-right axis is observed when stimulating NMs on the right-side of the head (N = 6 fish, n = 15 NMs, n = 760 neurons; Mann-Whitney U-stat = 1.5x105, p < 0.001). **D.** An illustration of a larval zebrafish indicating the position of individually stimulated NMs innervated by either the anterior lateral line (aLL; cyan) or posterior lateral line (pLL; magenta) on the left-side of the body. **E-F.** Contour heat maps in dorsal, sagittal, and coronal views represent the KDE of flow responsive hindbrain neurons (n = 2,504) that responded when stimulating individual NMs (N = 33 fish, n = 84 NMs; randomly sampled n = 10 NMs, bootstrapped n = 1k) from either the aLL or pLL on left-side of the body. **G.** Histogram and box-and-whisker plot of responsive neurons normalized by the number of NMs sampled in either the aLL (cyan) or pLL (magenta) along the left-right (F-stat = 0.12, p = 0.73, **i**), anterior-posterior (F-stat = 6.15, p = 0.01, **ii**), and dorsal-ventral (F-stat = 12.07, p < 0.001, **iii**) axes. **H.** Scatter plots with line of best fit of average coordinate position of flow responsive neurons along anterior-posterior (left; R^2^ = 0.01, F-stat = 0.11, p = 0.75) and dorsal-ventral (right; R^2^=0.45, F-stat = 8.93, p = 0.01) axes with respect to the relative position of each NM from head to tail. **I.** Heatmap representing KS-statistics for pairwise comparisons between aLL and pLL NMs coarse-grained distributions along the MON anterior-posterior axis (top) and dorsal-ventral axis (bottom). Lower KS values (blue) represent greater similarity, while higher values (red) represent more distinct distributions. **J.** Confusion matrices generated from a random forest model representing the moderate probability (65%) of successfully predicting the corresponding LL subtype (aLL = 67%; pLL = 64%; top) based on coarse-grained coordinates of the responsive neurons increased overall accuracy (84%) when predicting whether the stimulated NM was embedded on the head (83%) or tail (85%; bottom). **K.** A confusion matrix representing the probability (60%) of successfully predicting NM identity based on the coarse-grained coordinates of the responsive neurons from randomly selected NMs indicated a lack of spatial resolution to reliably differentiate responses across all sampled NMs. Scale bars represent 50 µm and error bars represent standard deviation of the mean.

In aquatic vertebrates, the spatial distribution of NMs along the anterior-posterior axis suggests that central processing of fluid motion may integrate sensory input according to the detection site. At the periphery, incipient somatotopy is evident in the primary projections of lateralis afferents onto the hindbrain^17^, where signals are segregated between the anterior and posterior LL ganglia. To determine whether the MON exhibits somatotopic organization of signals broadly distributed across the entire body, we stimulated individual NMs along the head and tail (**Figure 2D**). Stimulation of aLL NMs consistently activates neurons in the ventral MON, located within the prepontine and pontine regions (r1-3; **Figure 2E**). In contrast, stimulation of pLL NMs additionally activates a more dorsal and posterior region of the MON within the retropontine area (r4-6; Figure 2F).

The effects of LL subtype (i.e. aLL or pLL) and hindbrain recruitment patterns were assessed on normalized neuron counts along the lateral–medial, anterior-posterior, and ventral–dorsal axes of the MON. Along the medial–lateral axis, neither LL subtype (F_1,150_ = 0.02, p = 0.88), hindbrain region (F_1,150_ = 0.12, p = 0.73), nor their interaction (F_1,150_ = 0.23, p = 0.63) showed significant effects, indicating similar receptive field distributions across LL subtypes in both medial and lateral portions of the MON (Figure 2Gi). Along the anterior-posterior axis, there was a significant effect of hindbrain region (F_1,154_ = 6.15, p = 0.014), but not LL subtype (F_1,154_ = 0.02, p = 0.90) or interaction (F_1,154_ = 1.92, p = 0.17). Post-hoc pairwise comparisons (Holm-corrected) showed that aLL stimulation preferentially activated the anterior MON (p = 0.035; **Figure 2Gii**). Along the dorsal–ventral axis, a significant main effect of the hindbrain region was again observed (F_1,156_ = 12.07, p < 0.001), with more neurons recruited in the ventral MON during aLL stimulation. LL subtype (F_1,156_ = 0.02, p = 0.90) and the interaction (F_1,156_ = 1.13, p = 0.29) were not significant. Post-hoc testing confirmed enhanced ventral MON recruitment during aLL stimulation (p = 0.007; **Figure 2Giii**). Taken together, these results suggest that aLL stimulation preferentially recruits neurons within the anterior-ventral MON, whereas pLL receptive fields are more broadly distributed and overlapping with aLL receptive fields.

To further investigate the basis of this overlap, we examined response patterns of individual NMs (**Supp. Fig. 1**). Although we found no evidence of a continuous somatotopic map along the anterior–posterior axis of the hindbrain (R^2^ = 0.01, p = 0.75), we detected a significant gradient along the dorsal–ventral axis (R^2^ = 0.45, p = 0.01; **Figure 2H**). Specifically, the anteriormost NMs consistently activated the ventralmost MON neurons, while the posteriormost NMs preferentially recruited the dorsalmost MON neurons. To confirm receptive field distributions differed across NMs, we compared coarse-grained spatial coordinates of neuronal responses and found significant variation (KS-stat = 0.15, p < 0.001). Along the MON dorsal-ventral axis, pairwise comparisons revealed that pLL NMs located on the head had receptive field distributions similar to neighboring aLL NMs, whereas more distal pLL NMs on the tail showed distinct, non-overlapping patterns (**Figure 2I**). These results suggest functional overlap among head NMs, despite their separate ganglionic innervation prior to projecting to the MON. These findings support the idea that pLL NMs do not collectively function as a consolidated unit, but rather exhibit positional gradients in their receptive field topography depending on whether they are located on the head or tail.

To explore the organization of receptive field overlap at the single cell level, we identified MON neurons that responded to stimulation of multiple NMs. Coarse-grained neuronal coordinates (10 μm^3^) were filtered for voxels that contained responses to more than one NM within individual fish. The majority of neurons responded to only a single NM (n = 1,212 voxels), while a smaller fraction responded to multiple NMs (n = 108 voxels). These multi-responsive neurons typically responded to stimulation of 2–4 NMs, and co-activation was more common among spatially proximal NMs (**Supp. Figure 2A**). Very few neurons responded to both head and tail stimulation (n = 22 voxels; **Supp. Figure 2B**), suggesting functional segregation between these inputs.

Discrete somatotopic representation of head and tail was further tested using a random forest model, trained on ensembles of coarse-grained neuronal coordinates representing responses to each NM. The model achieved moderate accuracy in classifying whether the stimulated NM belonged to the aLL (67%) or pLL (64%), with an overall accuracy of 65% (**Figure 2J**). Predictive performance declined as spatial resolution decreased, with the minimum resolution sufficient for above-chance classification (∼50%) being 75 μm^3^ (**Supp. Fig. 3**). Interestingly, model performance improved when NMs were grouped by head or tail location, achieving 83% and 85% accuracy, respectively, suggesting a more salient somatotopic organization based on axial position rather than LL subtype. When predicting individual NMs based on coarse-grained neuronal coordinates, the random forest model achieved low accuracy (60%; **Figure 2K**), suggesting that while a general somatotopic organization exists between head and tail, significant overlap of receptive fields among neighboring NMs makes precise localization less distinct across the entire population.

For subsequent analyses, NMs were categorized as belonging to the head or tail based on their position relative to the animal’s center-of-mass, estimated at ∼1 mm from the rostrum within the swim bladder^36,37^. This anatomically informed division highlights a somatotopic organization in the MON that appears tuned to spatial location along the body axis relative to the center-of-mass, setting the stage for exploring how directional flow cues from the head and tail are distinctly represented.

### Sensory somatotopy within MON represents flow direction relative to center-of-mass

Until now, our analyses have focused on responses to anterior-to-posterior or “tailward” flow. To investigate how directional cues shape somatotopy in the hindbrain, we next examined responses to the reversed flow, referred to as posterior-to-anterior or “headward” (**Figure 3**). Hair cells are selectively sensitive to flow in a single direction which depends on the orientation of the ciliary bundle along the anterior-posterior axis. Hair cells of opposite polarities are antagonistically organized relative to one another and are equally represented within each NM^38^. Afferents can innervate multiple NMs but only synapse onto hair cells with matching polarity^22–24^. It is unknown whether opposing directional inputs project to distinct regions within the brain. Resolving this question is critical for understanding if the brain preserves directional specificity at early stages of processing, potentially shaping reflexive motor outputs tailored to directional cues.

**Figure 3.**
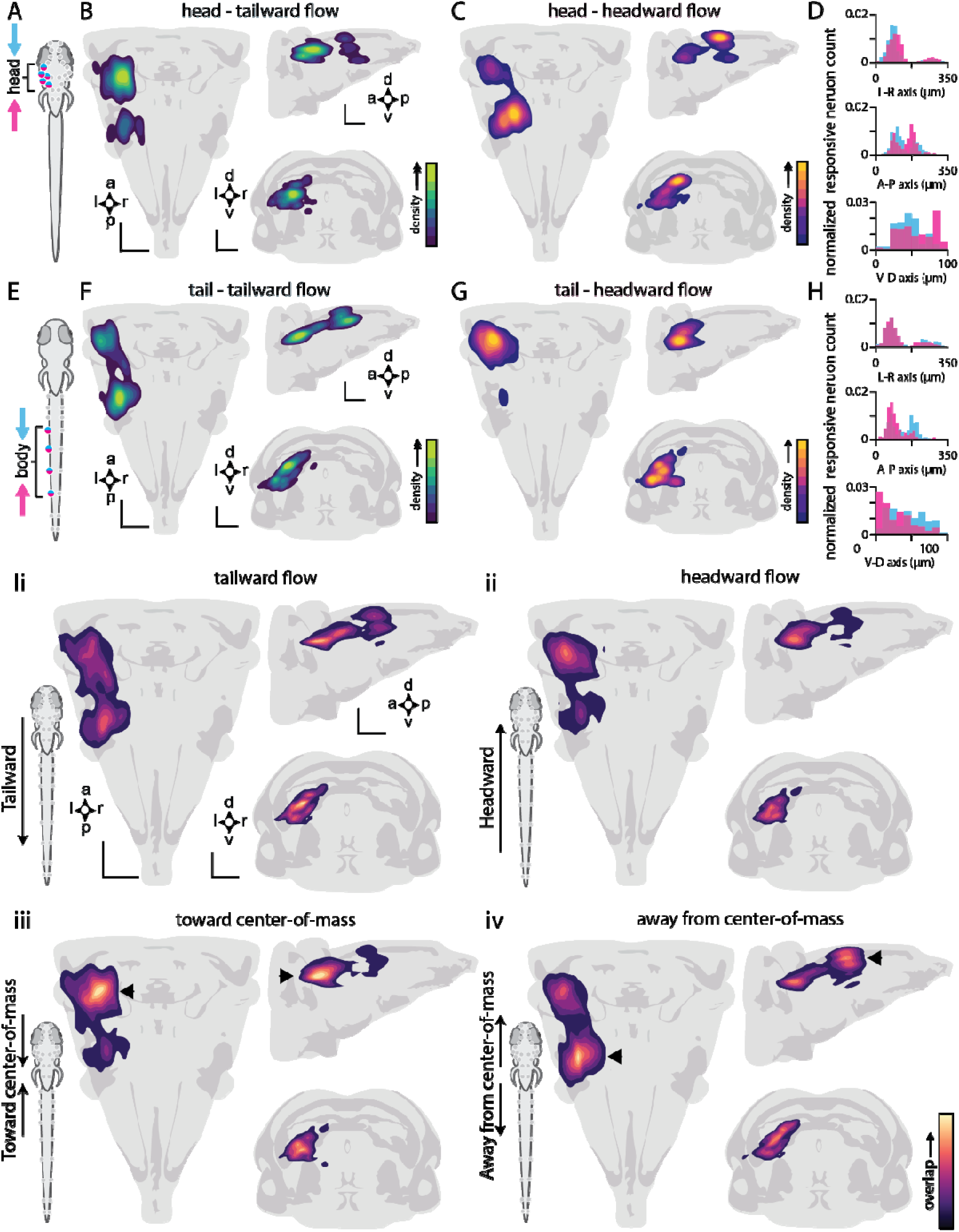
Receptive fields of head and tail change with respect to flow direction. **A**. An illustration of a larval zebrafish indicating the position of individually stimulated anterior lateral line (aLL) neuromasts (NMs) with both tailward (cyan) and headward (magenta) flow. Contour heatmaps indicate the relative kernel density estimates (KDE) of flow responsive neurons within the hindbrain in dorsal, sagittal, and coronal views when stimulating individual aLL NMs in the tailward direction (**B**) or headward direction (**C**). **D.** Histograms of normalized counts of hindbrain neurons that respond to stimulation of NMs located on the head of the animal with either tailward (cyan) or headward (magenta) flow (N = 11 fish, n = 16 NMs, n = 575 neurons). **E**. An illustration of a larval zebrafish indicating the position of posterior lateral line (pLL) NMs individually stimulated with both tailward (cyan) and headward (magenta) flow. Contour heat maps indicate the KDE of flow responsive neurons when stimulating individual pLL NMs in the tailward direction (**F**) or headward direction (**G**). **H.** Histograms of normalized counts of hindbrain neurons that respond to stimulation of NMs located on the body of the animal with either tailward (cyan) or headward (magenta) flow (N = 4 fish, n = 7 NMs, n = 663 neurons). **I**. Heatmaps of pairwise comparisons of MON activity using a three-dimensional KDE to determine the proportion of overlap of receptive fields encoding different flow directions along the head and body. Tailward flow demonstrated moderate overlap across the entire MON (42 ± 6%) with a maximum probability density function (pdf) overlap of 4.2 x 10^-6^ (**i**) and headward flow demonstrated low overall overlap (38 ± 6%) with a maximum pdf of 4.2 x 10^-6^ (**ii**). Flow toward or away from the center-of-mass demonstrated moderate overall overlap (44 ± 9% and 41 ± 6%, respectively) particularly in either the anterior-ventral (**iii**) or the posterior-dorsal (**iv**) MON with the maximum pdfs at 5.0 x 10^-6^ and 6.1 x 10^-6^, respectively (indicated by black arrows). Scale bars represent 50 µm.

To observe the recruitment patterns of MON neurons relative to stimulus direction, we investigated the recruitment of MON neurons during stimulation of NMs with bidirectional flow by again imaging 4-6 dpf *Tg(elavl3:H2B-GCaMP6f)* transgenic larvae (**Figure 3**). By positioning stimulus pipettes on either side of the target NM along the anterior-posterior axis, we deployed tailward flow followed by headward flow to volley an individual cupula back-and-forth to monitor hindbrain responses to both directions within the same NM and fish. When individual head NMs experienced tailward flow, we observed activity in the prepontine, pontine, and retropontine (rhombomeres r1 to r6) of the hindbrain (**Figure 3 A, B**), consistent with activation patterns previously observed (see **Figure 2E**).

Upon stimulation of the same head NMs deflected in the opposite direction (headward), we observed a dorsal portion of the MON activated more caudally, namely in the retropontine region (r4-6) with some overlap in the pontine region (**Figure 3C**). This pattern of activation was surprisingly similar to the previously described pattern of activation during tailward stimulation of tail NMs (see **Figure 2F**). Pairwise comparisons of distributions of tailward and headward flow responses on the head indicate anatomically distinct receptive fields along the anterior-posterior and dorsal-ventral axes within the MON (A-P axis: KS-stat = 0.25, p < 0.001; V-D axis: KS-stat = 0.29, p < 0.001; **Figure 3D**). When individual tail NMs experienced tailward flow (**Figure 3E**), we observed dorsal-caudal activation, but when reversing the flow to headward, we observed a significantly different distribution of responsive cells with a discrete rostral-ventral cluster emerging (A-P axis: KS-stat = 0.35, p < 0.001; V-D axis: KS-stat = 0.32, p < 0.001; **Figure 3F-H**). Strikingly, headward flow along the body generated a similar pattern of activation to activity associated with tailward flow along the head.

This unexpected observation motivated closer examination of overlapping receptive fields of flow direction relative to body position. When analyzing neuron ensembles across different flow directions, we observed overlap in the neural activity patterns. To quantify this effect, we fit a three-dimensional kernel density estimate (KDE) to the neural coordinates and computed the overlap coefficient to compare receptive fields between tailward and headward flow. Moderate overlap was observed across the entire MON during tailward stimulation of head and tail NMs (42%, **Figure 3I**), with a relatively low maximum probability density function (pdf) of 4.2 × 10^-6^. Headward flow exhibited better segregation between head and tail responses, reflected in a lower overlap of responsive neurons in the MON (38%, **Figure 3II**), while also maintaining a similar pdf maximum of 4.2 × 10^-6^. The low overlap within tailward and headward flow receptive fields across the entire body further supports a discrete somatotopic organization of stimulus direction across the head and tail.

When flow direction was analyzed relative to the center-of-mass, a distinct pattern of neural recruitment emerged. Flow directed toward the center-of-mass (tailward on the head and headward on the body) resulted in the greatest overlap in MON activation (44%, **Figure 3Iii)**, alongside an elevated maximum pdf of 5.0 × 10⁻□ (**Figure 3Iiii**). Similarly, flow away from the center-of-mass (headward on the head and tailward on the body) produced moderate overlap (41%) but showed the highest peak in overlapping activation (pdf max: 6.1 × 10⁻□, **Figure 3Iiv**). Considering flow relative to the center-of-mass rather than the whole body not only revealed slightly more overlap in receptive fields, but this overlap was concentrated in rostral or caudal regions of the MON (rMON or cMON, respectively).

Such a pattern suggested that spatially focused hotspots of activity dominate over broadly distributed overlapping sensory field maps. We therefore pooled responsive neurons according to directional cues relative to center-of-mass to reveal anatomically distinct sub-nuclei within the MON encoding flow toward or away from the center-of-mass (**Figure 4A**). We next analyzed the distribution of neurons within the newly defined subregions based on their respective NM’s position relative to the center-of-mass allowing for symmetric scaling of anterior-posterior positioning regardless of magnitude (**Figure 4Bi**). Effect size analyses revealed a robust anteroposterior separation of neurons within those that responded to flow toward the center-of-mass. Within the rostral MON, neurons responsive to head NM stimulation occurred more posteriorly while neurons responsive to tail NM stimulation were anteriorly located (Mann–Whitney p < 0.001, Cliff’s δ = -0.44). A complementary shift was observed dorsoventrally, with head NM responsive neurons located more dorsally than tail NM responsive neurons (Mann–Whitney p < 0.001, Cliff’s δ = 0.43). No meaningful spatial separation was observed laterally (Mann–Whitney p = 0.90, Cliff’s δ = 0.006; **Figure 4Bii**). Despite significant differences detected between the central tendencies of neuronal position in the receptive field encoding flow away from the center-of-mass, spatial segregation was less pronounced (dorsal-ventral: Cliff’s δ = 0.28, anterior-posterior: Cliff’s δ = -0.18) with negligible differences along the lateral axis (Cliff’s δ = 0.17; **Figure 4C**).

**Figure 4.**
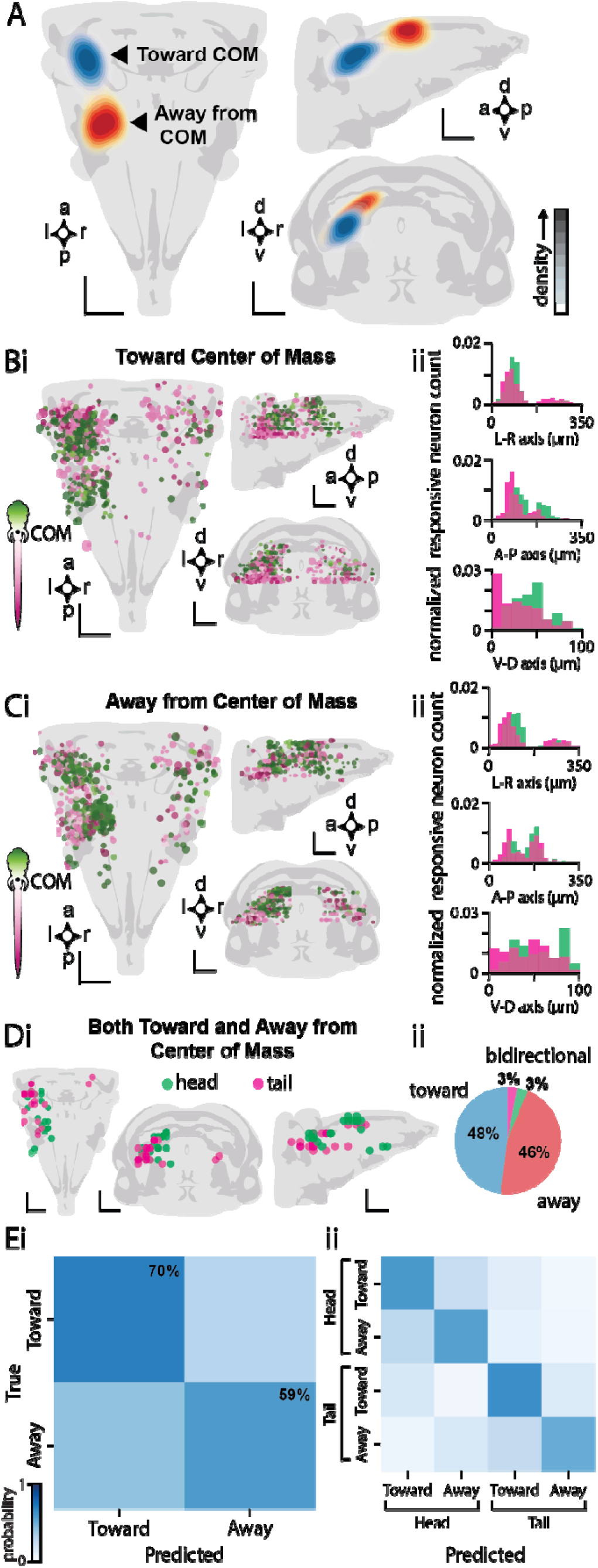
Flow responsive MON neurons discretely organize according to flow direction relative to the animal’s center-of-mass. **A**. Contour heatmaps indicate the relative kernel density estimates (KDE) of flow responsive neurons within the hindbrain in dorsal, sagittal, and coronal views reveal anatomically discrete receptive fields when stimulating individual NMs toward the center-of-mass (COM; blue, N = 12 fish, n = 20 NMs, n = 650 neurons) or away from the center-of-mass (red, N = 15 fish, n = 23 NMs, n = 588 neurons) **B.** Coordinates of individual responsive neurons to flow toward the center-of-mass color-coded to a normalized distance metric that symmetrically scaled values, enabling a direct comparison of anterior-posterior positioning regardless of magnitude. Dot diameter represents average response amplitude (0.1 - 1.0 ΔF/F) (**i**). Histograms of normalized responsive neurons counted across the left-right, anterior-posterior, and dorsal-ventral axes of the hindbrain (**ii**). **C.** Coordinates of responsive neurons to flow away from the center-of-mass (**i**). Histograms of normalized neuron counts across hindbrain axes (**ii**). **D.** Coarse-grained coordinates of responsive neurons to both flow toward and away from the center-of-mass in the same treatment revealing bidirectionally tuned neurons specific to head (green) or tail (magenta; **i**). Pie-chart demonstrates the low proportion of bidirectionally tuned neurons (6%) relative to those selective to flow toward (48%) or away from (46%) the center-of-mass (**ii**). **E.** A confusion matrix generated from a random forest model representing the moderate accuracy (65%) of successfully predicting whether the sensor experienced flow toward (70%) or away from (59%) the center-of-mass based on ensembles of coordinates of the responsive neurons during bidirectional stimulation of individual NMs (**i**). Predicting flow direction relative to the center-of-mass as well as whether it occurred on the head or tail modestly decreased predictability (58%; **ii**). Scale bars represent 50 µm.

We next identified neurons that responded to both flow toward and away from the center-of-mass to assess directionally nonselective responses. Bidirectionally tuned cells were defined as those activated by stimulation of the same NM under both flow conditions within a single larva, with responses detected in the same coarse-grained voxel (**Figure 4Di**). This approach revealed a subset of neurons that appear to encode the location of the stimulus (i.e. head or tail) irrespective of flow direction. These bidirectional neurons comprised a minority (6%) of flow-responsive neurons in the MON, compared to the majority that were directionally selective for flow either toward (48%) or away from (46%) the center-of-mass (**Figure 4Dii**). The small population of bidirectional neurons emphasizes the importance of the egocentric framework but may also suggest a parallel encoding strategy that supports location-based computations independent of flow direction.

To further test whether MON subregions encode both relative flow direction and axial position, we trained a random forest classifier on ensembles of coarse-grained neuronal coordinates representing responses to each NM under both flow directions. Model accuracy was low when attempting to classify individual NMs and flow direction relative to the center-of-mass (44%), indicating insufficient resolution in the MON map to reliably predict stimulus origin at the sensor level (**Supp. Figure 4**). Moderate accuracy was achieved when predicting whether the stimulated NM experienced flow towards (70%) or away (59%) from the center-of-mass, with an overall accuracy of 65% (**Figure 4Ei**). Predictive accuracy decreased when classifying head or tail during flow directed either toward (63% for head; 65% for tail) or away (55% for head; 45% for tail) from the center-of-mass, yielding an overall classification accuracy of 58% (**Figure 4Ei**). Rather than encoding axial location, the MON appears to prioritize egocentric directional information, effectively parsing ebbs and flows from the body’s center-of-mass, thereby simplifying the neural representation of distributed sensory input for downstream motor commands.

### Secondary projections from MON directly innervate reticulospinal neurons

To test whether these MON sub-nuclei contact motor command circuits, we investigated secondary projections from MON to command neurons, referred to as reticulospinal neurons. In fish and mice, numerous reticulospinal neurons express early in development the *vsx2* transcription factor^39^ and demonstrate functional specialization according to their date of birth^40^. While early-born *vsx2^+^* reticulospinal neurons drive escape and struggle behaviors, later-born *vsx2^+^* reticulospinal neurons are recruited for the spontaneous navigation composed of forward bouts or turns and deployed to explore and capture prey^40^. Population calcium imaging indicated that distinct subsets of *vsx2^+^* reticulospinal neurons are recruited to elicit forward versus turn bouts, and receive direct inputs from mesencephalic locomotor region neurons^41–43^. However, it is not clear whether MON neurons project onto specific sets of *vsx2*^+^ reticulospinal neurons to promptly select the motor output for either forward or ipsi- / contra-lateral directed turns.

To reveal the extent of rostral MON (rMON) connectivity onto *vsx2^+^* reticulospinal neurons we performed optical backfills of rMON soma by leveraging targeted multiphoton focusing on the pan-neuronal line expressing the photoactivatable GFP in combination with a red fluorescent protein labeling *vsx2^+^* neurons in the double transgenic *Tg(Cau.Tuba1:c3paGFP; vsx2:LOXP-DsRed-LOXP-GFP)*^a7437Tg;^ *^nns3Tg^* larvae (**Figure 5A**). The intense brightness generated at the photoactivation site made it challenging to reveal any ipsilateral axosomatic connectivity. However, this approach uncovered contralaterally projecting axons of rMON neurons contacting the soma of *vsx2^+^* reticulospinal neurons (**Figure 5A**). Putative axosomatic connections onto contralateral RoM3 cells in the rostral medulla responsible for driving forward locomotion were identified (**Figure 5Ai**) in addition to putative synapses onto MiD2i neurons in the retropontine responsible for forward locomotion and steering^44^ (**Figure 5Aii**). We also noticed posteriorly projecting secondary fibers running on ipsilateral and contralateral sides forming putative axo-dendritic synapses onto ipsilaterally-projecting *vsx2^+^* reticulospinal neurons (**Figure 5Aiii-iv**), previously found to be recruited during forward locomotion in the caudal medulla^40,43^.

**Figure 5.**
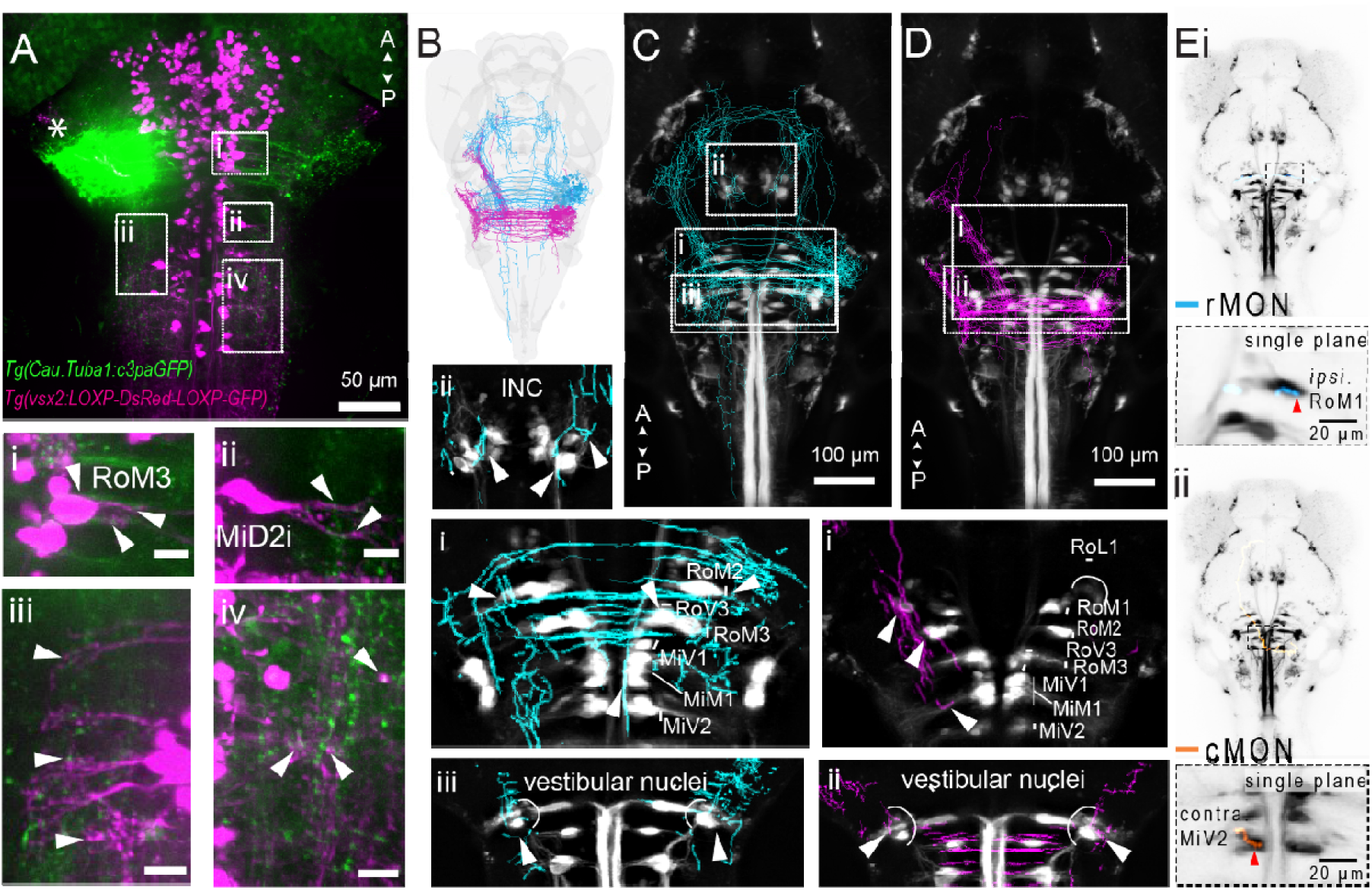
Anatomical connectivity of secondary projections from MON to motor commands referred to as reticulospinal neurons in the hindbrain, vestibular nuclei, interstitial nucleus of Cajal (INC) and torus semicircularis (inferior colliculus). **A**. Optical backfills of rostral MON soma (green) reveal extensive projections contralaterally to form putative connections (arrowheads) with *vsx2*^+^ (magenta) reticulospinal neuron soma such as (**i**) RoM3, (**ii**) MiD2i as well as lateral axo-dendritic connections to (**iii**) retropontine neurons ipsilaterally and (**iv**) rostrally medullary contralaterally (**iv**). Scale bars represent 10 µm and the asterisk marks the photo-activation site. **B**. Dorsal view of a whole brain with secondary projections from the MON color coded according to receptive fields encoding flow toward the center-of-mass (blue) or away from the center-of-mass (magenta) taken from the single-cell atlas MapZeBrain (https://mapzebrain.org/home). **C.** Dorsal view of rMON projections overlaid with reticulospinal neurons with 3 µm projections of planes that revealed extensive connectivity with reticulospinal neurons (arrows; **i**), innervation of the interstitial nucleus of Cajal (INC, **ii**), and bilateral connectivity with vestibular nuclei (**iii**). **D.** Dorsal view of cMON with unilateral projections onto canonical *vsx2*^+^ reticulospinal neurons (**i**) and bilateral projections to the vestibular nuclei (**ii**). Scale bar represents 100 µm **E**. Single cell traces of rMON (**i**) and cMON (**ii**) projections demonstrate unilateral axosomatic connections with reticulospinal neurons engaged during forward swimming and turns on either the ipsilateral or contralateral side.

To study how MON neurons project onto a broader population of reticulospinal neurons, we leveraged the virtual single-cell tracing of the publicly available brain atlas *mapZebrain*^45^ (https://mapzebrain.org/home) at the cellular resolution onto the canonical large caliber reticulospinal neurons named by Kimmel and colleagues in 1982^46^ (**Figure 5B**).

First, we found that rMON neurons (n = 43) exhibit an extensive connectivity profile forming robust, bilateral, putative connections onto large-caliber reticulospinal neurons distributed across both retropontine and pontine brainstem levels **(Figure 5C**). These reticulospinal targets include the major descending motor circuits that provide the primary drive for axial motor control (**Figure 5Ci**; arrowheads). The bilateral nature of rMON projections suggests a role in coordinating symmetrical motor output, which would be essential for forward locomotion as well as postural stability. Furthermore, rMON neurons also send projections to the interstitial nucleus of Cajal (INC) located in the diencephalon (**Figure 5Cii**). The INC, also referred to as the nucleus of the medio-lateral fasciculus^47,48^ (nMLF), serves as a critical integration center for the coordination of eye position, posture and locomotion^49–51^. In addition, the ventral commissure of rMON projections more often than the cMON reaches the torus semicircularis in the midbrain^29^ (**Figure 5C**). This projection positions rMON as a potential interface between locomotor command systems and higher-order feature detector circuits (i.e. torus semicircularis) homologous to the inferior colliculus in amniotes. In contrast, the caudal MON (cMON) neurons (n = 61) displayed a more selective and unilateral projection profile (**Figure 5D**). The dorsal commissure across the reticulospinal formation primarily terminates in a contralateral receptive field within the hindbrain^29^. The most prominent connections of cMON are observed onto the canonical reticulospinal neurons referred to as RoL1 and RoM2 (**Figure 5Di**). The cMON projections terminate on neuronal cell bodies, suggesting direct synaptic modulation of the spiking of reticulospinal neurons that encode forward locomotion during the optomotor response^44^. Some axodendritic connections were also observed, such as onto MiM1 which is also implicated in forward locomotion^44^. In addition to these forward locomotion targets, cMON neurons also project to reticulospinal neurons involved in turning, including RoM1, RoV3, and the MiVs^40,43,51–53^. Unlike the broader rMON projection pattern, cMON appears to project unilaterally to contact spatially restricted command neurons, particularly relevant for asymmetric steering responses. Notably, rMON and cMON share the characteristic of bilateral projections to the vestibular nucleus, indicating that both MON subdivisions contribute to vestibular system modulation (**Figure Ciii, Figure Dii**). This bilateral innervation pattern suggests a role in coordinating bilateral vestibular processing, which is essential for maintaining equilibrium during locomotor maneuvers and postural adjustments with direct relevance to the center-of-mass of the animal^54^. The unilateral projections from both the rMON and cMON provide the anatomical substrate for generating asymmetric motor commands necessary for directional steering behaviors. Axosomatic connections onto ipsilateral or contralateral reticulospinal neurons that are engaged during turning suggests the rMON and cMON contribute to steering either toward or away from the stimulus source, respectively (**Figure 5E**).

## Discussion

### Integration from single sensors spread over the entire body

Natural hydrodynamic environments are highly dynamic, shaped by currents and eddies that interact with the body at the fine scale of individual sensory loci. For larval zebrafish, these localized perturbations provide critical spatial and directional cues essential for survival. While flow may engulf the entire body, the lateral line (LL) system does not encode the absolute flow but the local changes in flow at the level of individual neuromasts^55^ (NMs). A fundamental question in sensory neuroscience is how the activation of spatially distributed sensors is compactly and reliably encoded in the brain.

The biological relevance of single-sensor inputs is well established in other sensory systems. For example, single mammalian photoreceptors in the retina can detect individual photons^56^, while single olfactory receptors can drive behavioral responses in mice^57^. These findings underscore that single sensory units, despite operating within a noisy, redundant system, can carry salient information that drastically shapes perception and behavior. In the LL, single NM deflection is sufficient to elicit escape responses^30^, emphasizing its behavioral importance and sensitivity. However, the downstream integration of this signal in the brainstem as a function of location of NM on the body and directionality of the flow had not been described.

By stimulating individual NMs, our results provide the most comprehensive anatomical and functional characterization of the MON in larval zebrafish to date. While it has been hypothesized that MON receptive fields could be organized by both somatotopic position and flow direction to enhance signal detection and noise robustness^58,59^, such organization has remained elusive— likely due to limited sampling and the confounding effects of bidirectional or bulk stimulation. By incorporating both aLL and pLL inputs, and systematically probing responses to flow in opposite directions, we reveal a spatially structured and directionally tuned map within the MON. This approach not only clarified how individual sensor responses are represented in the hindbrain, but also generated a composite view of MON activity that may inform how whole-body, multi-sensor input is processed during natural behaviors.

Previous studies, including brain-wide activity mapping^27^, have characterized responses to bulk flow across the trunk and tail but lacked the resolution to detect fine-scale topographic organization within the MON. Previous work using single NM stimulation has yielded mixed conclusions: bi-directional mechanical deflection identified directionally selective MON neurons^18^, whereas optogenetic activation of individual NMs found no evidence of somatotopy^28^. This is however expected considering that optogenetic methods stimulate hair cells of both polarities simultaneously, and thereby mask direction-selective responses. Moreover, up until now, the posterior ramus of the pLL has been exclusively probed when investigating central processing in larval zebrafish, representing fewer than half of total NMs and likely limiting detection of broader organizational principles along the entire animal. Contrary to the view of the MON as a homogeneous processing center, our findings reveal anatomically and functionally distinct subregions that differentially encode flow direction as a function of the sensor axial position. By sampling a greater diversity of NMs from both the aLL and pLL and parsing direction-specific responses, we demonstrate a structured somatotopic and directionally tuned organization. This expanded and refined view of MON topography provides new insights into how the brain integrates distributed sensor inputs into a coherent, egocentric representation of flow to generate the adequate motor output.

### Topographic flow mapping

Topographic maps are a hallmark of sensory systems, enabling precise localization and discrimination of stimuli based on spatial origin (Luo and Flanagan, 2007). In the LL, such somatotopy has been observed in the midbrain of other teleosts and amphibians^25,26,60–62^. Our findings suggest that a similar organization exists at the level of the hindbrain, where the MON exhibits discrete neural maps that integrate both spatial and directional cues into topographically organized receptive fields on each ipsilateral side. We also identified a small population of contralaterally responsive neurons, which are consistent with cross-midline projections thought to mediate lateral inhibition, helping to prevent bilateral co-activation and enhancing directional discrimination^63–65^. This lateralized organization is relevant for integrative reflexes such as rheotaxis, which require detection of gradients across the body midline to orient appropriately in flow^10^.

The somatotopy we observed along the MON dorsoventral axis during unidirectional tailward stimulation aligns with previous anatomical studies showing that pLL afferents project more dorsally than aLL afferents, which terminate ventrally^17,18,25,26,66^. However, resolving a clear somatotopic representation of the body’s anterior-posterior axis within the MON is more difficult due to overlapping termination fields of the aLL and pLL along this axis. Our single-sensor approach revealed substantial functional overlap between aLL and anterior pLL inputs, similar to observations in adult goldfish where some MON neurons responded to both head and rostral body stimulation, while other neurons responded selectively to caudal trunk and tail stimulation^66^. These results suggest that somatotopic organization in the MON reflects the positional origin of a sensor (e.g. head vs. tail) more than strict adherence to LL subtype (i.e. aLL vs pLL).

MON somatotopy is likely defined by the developmental emergence of primary projections from the periphery. In the pLL, early-born afferents from the first primordium innervate tail NMs and project dorsally onto the MON, while late-born pLL afferents from the second primordium target head NMs and terminate more ventrally^24,67,68^. Similarly, the aLL undergoes a mirrored developmental sequence along the anterior-posterior axis, where anterior NMs are innervated earlier, and posterior aLL NMs are targeted later^69^. However, aLL development occurs over a narrower window, potentially resulting in less stratification among the ventrally projecting aLL afferents innervating NMs sampled in this study. Therefore, the anterior-posterior MON topography we discovered is generally consistent with the temporal order of LL neurogenesis. It is difficult to determine the developmental origin of the directional neural map given no genetic tools exist to distinguish between polarized afferents and their anatomical projections. Prior to our findings, there was no evidence that polarized lateralis afferents transmitting opposing directional information terminated onto different groups of MON neurons in the hindbrain.

### Differential connectivity onto motor command neurons and sensorimotor integration centers

Our findings support two differential patterns of projections along the antero-posterior axis of the MON^29^. The differential projection patterns of rMON and cMON suggest complementary yet distinct functional roles within the broader context of motor control and postural regulation. The different projections patterns suggest distinct pathways of motor control with short latency recruitment at the level of the reticulospinal neurons and longer timescale processing at the level of the diencephalon. The organization of these connections provides insight into how MON neurons collectively orchestrate complex locomotor behaviors through their influence on reticulospinal pathways.

The bilateral projection pattern characteristic of rMON neurons likely facilitates forward swimming behaviors through the generation of symmetrical motor commands. By simultaneously activating reticulospinal neurons on both sides of the brainstem, rMON can coordinate the rhythmic, bilateral muscle contractions necessary for propulsive locomotion. This symmetrical activation pattern would produce the coordinated undulatory movements typical of forward swimming, where left-right motor output remains balanced and phase-locked. Conversely, the predominantly unilateral projections of cMON neurons provide the anatomical framework for generating turning behaviors and directional changes. Asymmetrical activation of reticulospinal pathways through unilateral cMON influence would create differential motor output between the left and right sides of the body, resulting in steering maneuvers.

The unique rMON projections to the interstitial nucleus of Cajal further expand the postural regulatory capacity of the MON system. The INC plays a crucial role in controlling head positioning, gaze stabilization, and postural maintenance through its connections with oculomotor and neck motor systems^48–51^. Through this diencephalic connection, rMON neurons encoding flow toward the center-of-mass may influence higher-order postural control mechanisms that coordinate head-body relationships and maintain appropriate spatial orientation during locomotor activities. This connection suggests that rMON may serve as a critical link between locomotor circuits and postural control systems, enabling integrated control of whole-body movement and posture.

The center-of-mass serves as a universal reference that allows the integration of sensory inputs from different modalities. For example, vestibular inputs provide information about body tilt, and the LL encodes spatial and directional information about flow. If both are encoded relative to the center-of-mass, the brain can more easily combine these signals to generate appropriate movement responses. In fact, the MON functions as the sensorimotor integration center for both systems and we discovered bilateral projections to the vestibular nucleus from both cMON and rMON. MON neurons likely modulate vestibular processing and, consequently, postural reflexes and spatial orientation mechanisms. This vestibular influence would be particularly important during dynamic locomotor behaviors where continuous postural adjustments are required to maintain stability and orientation.

Future studies should aim to characterize the recruitment patterns of the reticulospinal neurons at the population level depending on stimulus direction, sensor position, and stimulus amplitude. Uncovering the stimulus parameters that lead to transitions between forward locomotion and steering or ipsilateral and contralateral recruitment will reveal fundamental principles about sensory mediated action selection.

### Egocentric directional system

The center-of-mass provides a stable, intrinsic frame of reference that is invariant to body movement and crucial for processing motor control in a way that supports effective navigation and environmental interaction. A key insight from our study is that directional inputs to the MON are organized relative to this egocentric reference frame reflecting a transformation from raw peripheral input to a spatially meaningful codex. Though individual LL afferents themselves do not independently encode flow direction^55^ the prevalence of mono-directionally tuned neurons in the hindbrain, especially compared to higher-order brain regions, suggests that the MON is the major site for directional processing^27^. While our study did not sample dorsal pLL NMs, their polarization along the dorsal-ventral axis and anatomical position above the center-of-mass^21^ suggest that dorsal pLL NMs may also conform to this egocentric framework by encoding vertical flow toward or away from the body core. Such an organization could reduce reliance on additional computational mechanisms like intrinsic coincidence detectors that compare input across multiple afferents^55^ or inhibitory–excitatory ensembles tuned to opposing directions^10,18^. By aligning directional tuning with an egocentric map of sensor position, the MON appears optimized to extract behaviorally relevant information about fluid motion relative to the body, rather than its absolute spatial source.

The question then remains, how does the MON encode directional information from biologically relevant flow stimuli from the environment to inform action selection? While strong flow cues can activate fast escape circuits via Mauthner neurons, bypassing the MON^70^, more subtle cues must first be processed in the MON to guide more directed behavioral outputs. For instance, predator approaches and strikes generate bow waves that propagate flow toward the center-of-mass of the prey larval zebrafish, which must be integrated to command contralateral turns that facilitate avoidance maneuvers^16,71,72^. Alternatively, laminar flow directed away from the center-of-mass along the tail biases for ipsilateral turns to steer and maintain counter-current orientation during rheotaxis^9,10^. It is important to maintain that stimulus magnitude plays a critical role in behavioral responses and our experimental paradigm intentionally falls within a low amplitude flow regime. While external flow cues drive context-specific responses through MON processing, the same egocentric framework may also support the interpretation of self-generated fluid motion during locomotion.

As animals move through their environment, their own body movements also generate fluid disturbances that impinge upon the LL. In adult fish, populations of MON neurons often respond transiently to this reafferent flow, rapidly adapting to rhythmic self-generated signals during swimming or respiration^73–76^. The degree to which larval zebrafish experience and process such flow depends critically on the scale and mechanics of their locomotion. At their small body size (< 5 mm), low-amplitude forward swimming (< 0.3 ms^-1^) produces relatively symmetric near-field flows mirrored across the left-right axis, with minimal net vorticity^77^ due to dominate viscous forces^78^ (*Re* < 300) that likely prevent NM deflection and subsequent recruitment of MON neurons. Despite this, it is possible that MON neurons respond to sudden changes in locomotor output such as the preparatory stroke to execute high amplitude turns that generate large asymmetries in the near-field vorticity along the trunk and tail of the animal^77^. These movements increase the inertial forces^78^ (*Re* ≈ 1500), raising the likelihood that NMs are deflected. Importantly, such asymmetries would result in directionally biased flow fields relative to the center-of-mass, selectively activating MON subregions tuned to motion away from or toward the center-of-mass depending on the curvature and direction of body movement.

This suggests that egocentric directional encoding in the MON may be critical not only for interpreting environmental stimuli but also for proprioceptive sensing during locomotion. By aligning directional sensitivity with the center-of-mass, MON neurons may provide a discretized map of fluid disturbances that reflect dynamic changes in posture and trajectory along the anterior-posterior and left-right axes. Corollary discharge mechanisms suppress predictable components of this self-generated flow at the periphery^73,79–81^, but at peak body curvature the residual directional input is likely passed on to the MON^32,76^. Within this narrow temporal window, directionally tuned MON neurons could act as encoders of body curvature during accelerations, integrating flow direction signals in an egocentric reference frame. Additionally, a summation of population level activity of the MON across long timescales could ostensibly capture fluctuations in reafferent feedback to reflect changes in navigational strategy such as wandering or cruising^82^.

## Materials and Methods

### Animals and transgenic lines

Animal handling and procedures were validated by the Paris Brain Institute (ICM) and the French National Ethics Committee (Comité National de Réflexion Éthique sur l’Expérimentation Animale; APAFIS no. 2018071217081175) in agreement with European Union legislation. To avoid pigmentation, all experiments were performed on *Danio rerio* larvae of AB background with the *mitfa^−/−^* mutation. Adult zebrafish were reared at a maximum density of eight animals per liter in a 14/10□h light–dark cycle environment at 28.5□°C. Larval zebrafish were typically raised in Petri dishes filled with system water under the same conditions in terms of temperature and lighting as for adults. Transgenic larvae expressing PA-RFP and used for the photoconversion experiments were exceptionally raised in complete darkness to minimize global photoconversion at the early stages. Experiments were performed at 20□°C on animals aged between 4 and 6 dpf. The following transgenic lines were used: *Tg(elavl3:H2B-GCaMP6f)^jf7Tg^* ^33^ & *Tg(Cau.Tuba1:c3paGFP)*^a7437Tg^; *Tg(vsx2:LOXP-DsRed-LOXP-GFP)^nns3Tg^*.

### Neuromast stimulation

Larvae were immobilized by immersion in a paralytic (20 μL of 1 mg/mL α-bungarotoxin [Tocris]) in E3^83^. Larvae were transferred to a coverslip bottomed recording dish with a well containing a semi-circle silicone elastomer (Sylgard 184). Larvae were orientated dorsal-side up and then pinned in two locations through the notochord (i.e. above the anus and the tip of the tail) on the left-side of the animal and into the diameter segment of the semi-circle elastomer bed using etched tungsten pins (<50 μm diameter; **Figure 1A**). A third pin was inserted below the mandible to level and stabilize the head for imaging. Anesthetics such as MS-222 (Tricaine) and the low temperature melting agarose could not be used to immobilize or encapsulate the animal due to their deleterious effect on the delicate cupula that is essential in the flow detection process^84,85^. Blood flow was monitored as a metric of animal health and only trials with robust blood flow throughout the experiment were considered for analysis. A stimulus pipette tip (diameter: ∼10 μm), fabricated from borosilicate glass capillaries (World Precision Instruments, Inc.; 1B150F-4) pulled on a model P-1000 Flaming/Brown micropipette puller (Sutter Instruments, Novato, CA), was filled with bath solution (E3) and aligned along the anterior-posterior axis approximately 20 μm anterior (unless otherwise specified) to the neuromast. The stimulus pipette was connected to a pneumatic microinjector (PV 820 Pneumatic PicoPump, WPI) which applied pressure to eject fluid from the pipette for 5 s, preceded and followed by a 20 s pre- and post-stim period, respectively. The stimulus program (Clampex 10.1) was controlled via a low-noise digitizer (Digidata 1440A). The effect of the controlled flow stimuli on cupula deflection was captured using an upright confocal spinning disk microscope (Yokogawa CSU-W1; Examiner Z1; LaserStack), and images were acquired at 1 frame/s with a field of view (FOV) of 680 × 680 μm (pixel size = 0.33 μm) using a X40 NA = 1.0 objective and using Slidebook v.2024.1 software. Kinocilia height (μm), tip deflection distance (μm), and deflection angle were all quantified using ImageJ (1.54f, NIH, Bethesda USA). Proximal, off-target neighboring neuromasts were also monitored to ensure the gentle fluid jet exclusively deflected the target neuromast. After image acquisition at one neuromast, the stimulus pipette was repositioned at a different neuromast to repeat the experiment.

### Calcium imaging

We examined the responses of hindbrain neurons to the mechanical stimulation of individual neuromasts along body axes. Larvae expressed the nucleus-localized fluorescent calcium indicator H2B-GCaMP6f with a pan-neuronal promoter *(Tg(elavl3:H2B-GCaMP6f)^jf7Tg^* ^33^. The stimulus protocol during calcium imaging ejected fluid from the stimulus pipette for 5 s, and the rest between stimuli was extended to 40 s. The protocol elicited three sweeps of this fluid puff and rest sequence. The first sweep was preceded by, and the last sweep was followed by a 20 s period of inactivity. Displacement of the kinocilia tips was confirmed by manually triggering the pneumatic transducer prior to repositioning the FOV for calcium imaging in the hindbrain. Changes in fluorescence were recorded at a single plane within the hindbrain using an upright confocal spinning microscope (CSU-X; Examiner Z1; LaserStack). Images were acquired using a X20 NA□=□1.0 objective, using Slidebook v.6.0. Spinning disk time series were acquired at 5 frames/s with a FOV of 340□×□340□μm (pixel size□=□0.66□μm). We systematically repeated the stimulus protocol while imaging at increasing depths separated by 10 μm (≤10 planes) to create a partial Z-stack volume of the hindbrain during neuromast stimulation. After calcium imaging at multiple planes, the stimulus pipette was repositioned at a different neuromast, and the protocol was repeated. We systematically recorded an anatomical Z-stack (100 planes separated by 1 μm) of the hindbrain before and after imaging calcium activity in each plane for reference during brain registration.

We investigated which hindbrain neurons could be responsive to specific stimulus parameters such as flow direction. To test if calcium activity patterns differed depending on flow direction, two identical stimulus pipettes were positioned along the anterior-posterior axis equidistantly on either side of an individual neuromast. The stimulus protocol was performed to deflect the kinocilia in the anterior-posterior direction, and then repeated while puffing from the twin pipette to deflect in the posterior-anterior direction.

Analysis of functional hindbrain imaging began with extraction of ROIs from the imaged volume from each trial (e.g. NM) of each fish. First, the time-series for each plane of each Z-stack was manually registered to a reference brain atlas so that approximate neuron position could be compared across individuals (ImageJ). Then, 2-step registration accounted for potential motion in 2D, and ROIs of all cells with their respective ΔF/F(t)□=□(F(t)□−□F0)□/□F0, where F(t) is the fluorescence at time t and F0 is the cell baseline fluorescence (median fluorescence value of 20 s period pre-stimulus), were automatically obtained in an open-source functional segmentation pipeline^86^ (Suite2p). Using custom-written Python script, we determined if each ROI was a flow responsive neuron if it met the criteria of ΔF/F(t) exceeding 4x sigma of F0 (defined independently for each trial), exceeding a minimum slope (0.025), and an accurate fit of the decay time constant (*τ*) during all three stimulus periods. Within each response interval, the decay time constant (*τ*) was estimated by fitting a mono-exponential decay model to the falling phase of the fluorescence response, starting from its peak: *f*(*t*) = *A*⋅*e*^−*t*/*τ*^+*C*, where *A* is the amplitude, *τ* is the decay constant, and c is an offset. The model was fitted using nonlinear least squares optimization (Levenberg–Marquardt algorithm), with initial parameter guesses set as *A* = peak − baseline, *τ* = 1.5, and *C* = baseline. Fits were also excluded if the peak occurred in the first frame of the stimulus window, or if the fitted τ exceeded a ceiling of 20 s. In addition, only ROIs with *τ* that shared 95% similarity across all three stimulus periods were included. With this approach, we aimed to conservatively give an overview of how many hindbrain neurons were consistently responsive to single neuromast stimulation.

### Lateral line ablation

We verified that the stimulus paradigm was selective to lateral line stimulation by performing chemical ablations. Calcium imaging of hindbrain activity was first recorded during stimulation under control conditions (E3). Then the membrane impermeant calcium chelator BAPTA (ThermoFisher Scientific, invitrogen) was bath applied at 5mM and incubated for 20 min to disrupt tip-links between stereocilia necessary for mechanotransduction^35^ and calcium imaging of hindbrain activity was recorded again while stimulating the same neuromast. To assess whether the number of responsive neurons was significantly higher in the control conditions compared to BAPTA conditions within individuals, we first tested for normality using the Shapiro-Wilk test followed by a paired one-tailed Student’s t-test.

### Spatial density estimation of MON

To quantify the spatial distribution of responsive neurons within the hindbrain across different neuromasts and flow directions, we employed a kernel density estimation (KDE)-based bootstrapping approach (n = 1000 iterations) to reduce biases introduced by oversampling of specific NM identities and individual variability. We first filtered the data based on experimental conditions (e.g. NM identity, NM position relative to center-of-mass, flow direction, etc.). For each bootstrap iteration, we randomly sampled a unique combination of individual, NM, and flow direction (without replacement). When relevant to restrict analysis to ipsilateral activity, coordinates were excluded beyond the midline. For each anatomical view (i.e. dorsal, sagittal, and coronal), a two-dimensional KDE was computed using a Gaussian kernel with a fixed bandwidth scaling factor (0.25). KDEs were visualized as contour maps, with a consistent number of contour levels (n = 10) and a defined minimum density threshold to facilitate comparison across anatomical view and conditions. To visualize the spatial extent of neuronal distributions of the left MON in 3D, we generated volumetric masks based on KDE of responsive neuron coordinates from ipsilateral NMs. The bootstrapped 3D coordinates from each anatomical view were used to compute a three-dimensional KDE. The KDE was evaluated on a uniform voxel grid (100×100×100) spanning the minimum and maximum coordinates in each axis. The KDE bandwidth was scaled by a factor of 0.25, consistent with earlier 2D contour visualizations. A density threshold was then set at the 95th percentile of the computed 3D density volume to isolate regions of highest neuronal concentration and generate a polygonal mesh as an isosurface representing the outer boundary of the MON (Lorensen and Cline 1988).

### Classifier-based spatial mapping

To investigate the spatial encoding of NMs or LL subtype, the three-dimensional coordinates of responsive neurons were aggregated for each individual NM which then served as input features for a supervised classification model. To account for spatial uncertainty introduced during registration to a common brain atlas, the coordinates were coarse-grained (10 µm³) to account for any margin of error around each neuron position. This discretization prevents the model from overfitting to spurious precision and reflects the biological variability and registration imprecision inherent in the dataset. For classification, we implemented a random forest model^87^ to distinguish between individual NMs, LL subtype, and axial position (i.e. head vs tail). For each NM sampled all associated neurons that responded to the same NM stimulation in a given individual were included. To investigate spatial encoding of directional movement relative to the center-of-mass, we trained another random forest classifier to distinguish coarse-grained (1□µm³) neuronal coordinates associated with stimuli either toward or away from the center-of-mass. To assess finer distinctions in directional somatotopy, we expanded the classification scheme to include four classes incorporating both anterior–posterior anatomical identity (i.e. head vs tail) and the directional alignment with respect to the center-of-mass (i.e. toward or away). To explore sensor-specific contributions to directional processing, we then trained a random forest model to classify neurons into distinct categories for each unique NM subtype and flow direction. Across models, the datasets were filtered to include only individuals with a minimum of three responsive neurons corresponding to each individual NM to ensure stability in label distribution. The dataset was split into training and testing subsets (80/20) using stratified sampling to preserve class proportions and then trained on the aggregates of 3D neuronal coordinates associated with a given NM, flow direction, and individual. The classifier was trained with the following parameters: 50 trees, a maximum tree depth of 20, and a minimum of 10 samples required to split an internal node. Model performance was assessed using a confusion matrix and accuracy score. Feature importances were extracted to quantify the relative contribution of each spatial axis to the classification.

### Quantifying overlap of receptive fields

To assess the degree of spatial overlap between receptive fields for distinct directional flow conditions in the MON, we computed a 3D overlap coefficient (OVL) based on KDEs of flow responsive neuron distributions. A random aggregate of responsive neuron coordinates associated with a unique combination of NM type, flow direction, and individual was sampled with replacement and bootstrapped (n = 1,000) to account for variability in sampling and across fish or NMs. All ROIs corresponding to the sampled combination were retained. The dataset was spatially restricted to only include responses ipsilateral to the stimulated NM. For each condition pair (i.e. flow toward center-of-mass, flow away from center-of-mass, tailward flow, and headward flow), we fit separate KDEs in 3D space to evaluate the minimum to maximum extent of both distributions. The overlap coefficient was computed as the integral of the minimum value (min) between the two KDEs (f1 and f2) at each point (x,y,z):

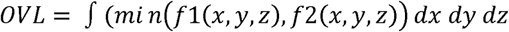

The OVL value ranges from 0 (no spatial overlap) to 1 (identical spatial distributions), and serves as a quantitative measure of similarity between the two receptive fields in the MON. To assess the robustness of spatial similarity estimates, we applied an additional bootstrap resampling procedure (n = 100 iterations) for each condition pair. For each bootstrap, neuronal coordinates from both conditions were resampled with replacement, and the OVL was recalculated. The resulting distribution of OVL values was used to compute the mean, 95% confidence interval (CI), and a similarity rating. To evaluate whether topographical representation of sensor position relative to the center-of-mass occurred within the flow toward or flow away receptive fields, we analyzed the spatial distribution of flow responsive neurons within each condition. To assess whether MON neurons corresponding to NMs located either anterior or posterior to the center-of-mass were spatially segregated depending on the flow direction, we conducted KS-test to detect differences in the spatial distributions, Mann-Whitney U test to assess median differences, and Cliff’s δ to capture non-parametric effect sizes. Together, these tests provided complementary insight into both the significance and magnitude of somatotopy representing opposing flow-responsive populations within NM populations anterior or posterior to the center-of-mass.

### Optical backfills

*Tg(Cau.Tuba1:c3paGFP)* with *Tg(vsx2:LOXP-DsRed-LOXP-GFP)^nns3Tg^* larvae were raised in darkness to avoid unintended exposure to near UV-light and subsequent photoactivation of c3paGFP. At 4 dpf, larvae were anesthetized with tricaine (MS-222, 0.02%, Sigma-Aldrich) and embedded dorsal-side up in 3% low melting-point agarose prepared in aCSF in a 35-mm petri dish. Photoactivation was performed on a multiphoton microscope (2p, Intelligent Imaging Innovations) where we scanned the rostral MON (50 μm depth) at 800 nm with a 8 μs dwell time in each plane (1 μm), for ∼5 min. After the procedure, the dish was filled with aCSF, the larvae were unmounted and allowed to recover overnight. Larvae were then anesthetized and mounted in agarose again for imaging of optically backfilled neurons. The retrogradely labeled neurons were imaged on a confocal spinning-disk microscope as described above.

### Single Cell Tracing

To investigate the single-cell tracing library of the mapZebrain atlas (https://mapzebrain.org/home), two regions of interest (ROIs) were defined to encompass the caudal and rostral MON. All neurons with cell bodies located within these ROIs were extracted, and the resulting image files were analyzed using Fiji (https://fiji.sc/Fiji) software with the Simple Neurite Tracer (STN) plugin for morphological quantification^88^.

## Acknowledgments

We would like to thank the Wyart lab “SIBBIL” (https://wyartlab.org; https://parisbraininstitute.org/paris-brain-institute-research-teams/sibbil-navigation-sensorimotor-integration-brain-body-integration-lab) for providing inputs along the way - particularly M. Dhanasekar, G. Sridhar and A. Carlos Costas for constructive discussions that helped shape the manuscript. We thank S. Nunes Figueiredo, M. Dicu, A. Arneau, N. Jezequel, C. Lejeune, B. Daboval of the core facility Pheno-Zfish for fish care. We thank F. Engert, M. Ahrens, S.-I. Higashijima for sharing published transgenic lines. E.L. was supported by the Human Frontier Science Program (HFSP-LFP). M.C.T. was supported by the Campus France PRESTIGE postdoctoral research fellowship (2017-2-0035). This project benefited from funding from a 2020 Fondation Bettencourt-Schueller (FBS-don-0031) award (Identity and organization or neuronal networks controlling exploration), a New York Stem Cell Foundation (NYSCF) Robertson Award 2016 grant 332 (NYSCF-R-NI39), the 2020 Prize Equipe ‘Fondation pour la Recherche Médicale’ (FRM-EQU202003010612) ‘Neuronal circuits underlying navigation: from genes to behavioral models’, a 2020 European Research Council Consolidator grant no. 101002870 (2021–2026) ‘Exploratome: Circuit mechanisms underlying sensory-evoked navigation’ and a National Institutes of Health grant no. 1U19NS104653-01 awarded to C.W., the European Union’s Horizon 2020 research and innovation programme under a Marie Skłodowska-Curie grant no. 813457 awarded to C.W. The project benefited as well from the support of the Agence Nationale pour la Recherche (ANR) ANR-22-CE37-0023 LOCOCONNECT, ANR-23-CE16-0017-02 RocSMAP, ANR-24-CE16-7992 CIRCOLOCO, ANR-21-CE14-0042 MOTOMYO and ANR-21-CE13-0008 ASCENTS.

---

**Supplementary Figure 1.**
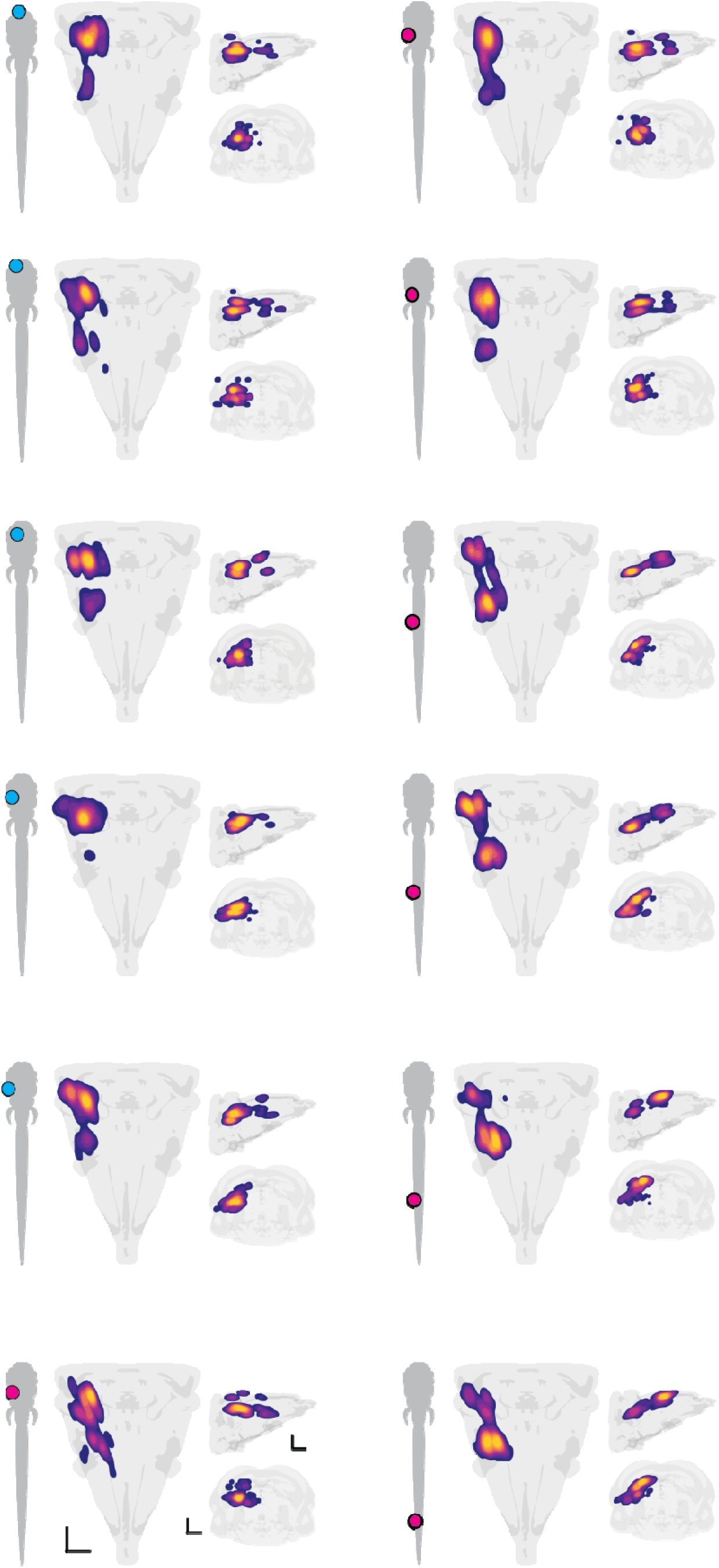
MON activity for each neuromast along the anterior-posterior axis. Contour heat maps indicate the KDE of flow responsive neurons within the hindbrain in dorsal, sagittal, and coronal views when stimulating individual neuromasts (NMs). Dots on the larval zebrafish silhouettes indicate the relative position of aLL (cyan) and pLL (magenta) NMs corresponding to each heatmap. Scale bars equal to 50 μm.

**Supplementary Figure 2.**
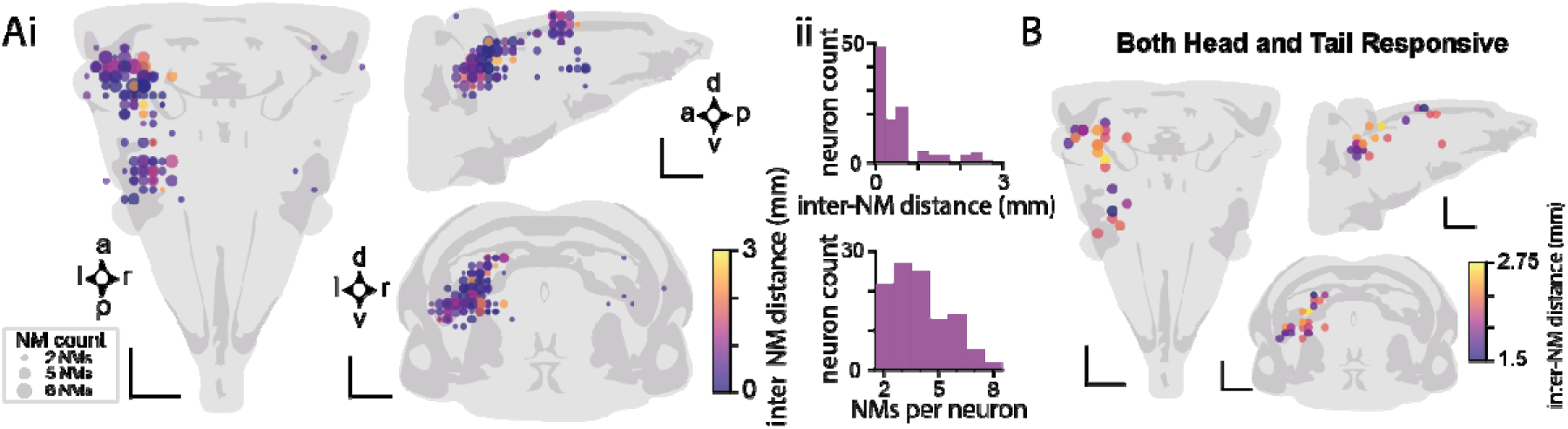
Neurons responsive to multiple neuromasts. **A.** Coarse-grained coordinates of neurons that responded to multiple neuromasts in a given individual. Color gradient represents the distance between two neuromasts while dot size represents the number of unique neuromasts that led to responses in that neuron. Coordinate plots are represented in dorsal, sagittal, and coronal views (**i**). Histograms indicating proximally located NMs more frequently led to responses in the same neurons (**ii**, top) and that those neurons that did respond to multiple neuromasts were typically sensitive to 2-4 (bottom). **B.** Coarse-grained coordinates of neurons that responded to both head and tail neuromast stimulation in a given individual. Color gradient represents the distance between the two neuromasts. Scale bars represent 50 μm.

**Supplementary Figure 3.**
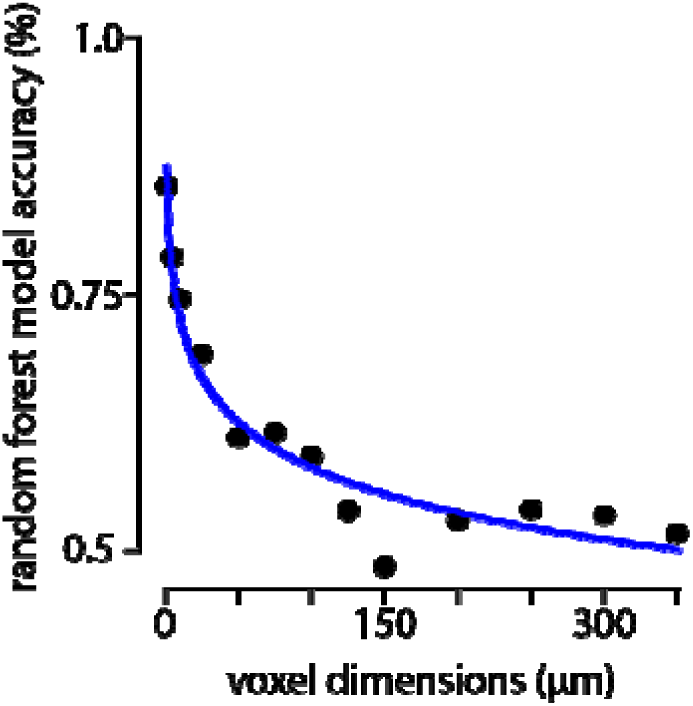
Relationship between coarse-graining and accuracy of random forest model. The overall accuracy of the random forest model predicting the corresponding LL (i.e. aLL or pLL) based on different levels of coarse-graining represented by increasing voxel dimensions (x, y, z) containing coordinates of the responsive neurons from randomly selected NMs. The blue curve represents a fitted logarithmic trend indicating accuracy decreases as voxel size increases, with accuracy decreasing below 60% at voxel sizes greater than 75 x 75 x 75 μm.

**Supplementary Figure 4.**
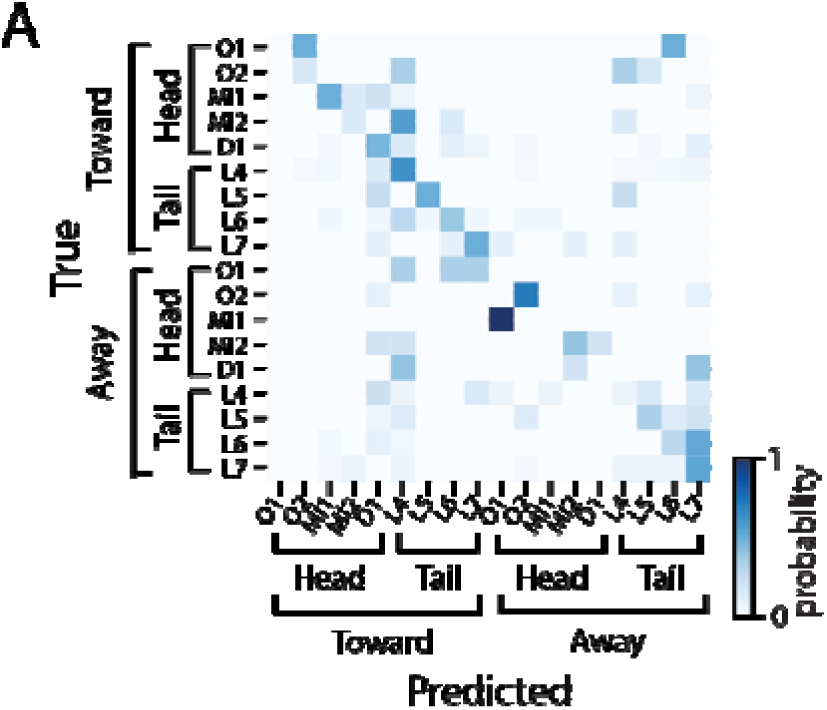
Confusion matrix for each neuromast and flow direction. A confusion matrix generated from a random forest model representing the low probability (44%) of successfully predicting NM identity based on the coordinates of the responsive neurons from individual NMs indicated a lack of spatial resolution to reliably differentiate stimulus direction and NM position across all sampled NMs.

