## Supplementary figures and images for "Whole-body central processing of lateral line inputs encodes flow direction relative to the center-of-mass"

### Supplemental Figure 1

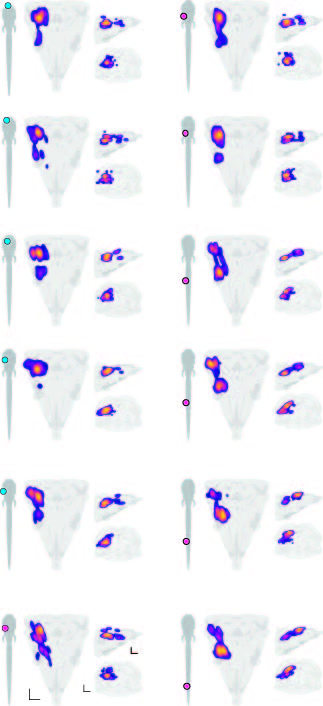

### Supplemental Figure 2

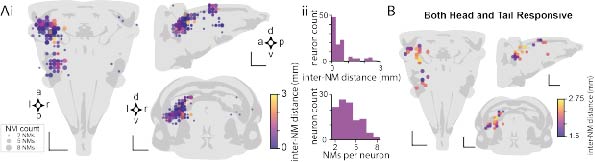

### Supplemental Figure 3

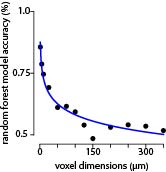

### Supplemental Figure 4

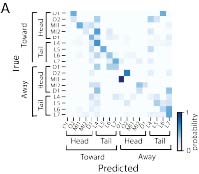
